# HaloPROTAC-mediated degradation reveals SOX10-dependent gene expression changes in melanoma cells

**DOI:** 10.64898/2026.10.19.760213

**Authors:** Ashleigh L. Purvis, Christine M. DeRosa, Andreas P. Pintado-Urbanc, Khushi Agnish, Kevin Jiang, Antonella Bacchiocchi, Ruth Halaban, Sarah A. Slavoff

**Author notes:** To whom correspondence should be addressed: Sarah Slavoff.

## Abstract

The transcription factor SOX10 is required for melanoma cell proliferation. However, SOX10 expression in single cells is heterogeneous, and low SOX10-expressing melanoma cells exhibit a dormant-invasive phenotype associated with drug resistance and metastasis. Despite longstanding interest in SOX10 as a diagnostic marker and potential therapeutic target in melanoma, the function and targets of SOX10 remain incompletely understood. New tools for the study of SOX10 are needed to address these gaps in our knowledge. In this study, we engineered a patient-derived WW165 melanoma cell line for programmable SOX10 targeted proteolysis. We achieved this by replacing the endogenous copies of *SOX10* with a SOX10-HaloTag transgene that supports cell growth, but can be rapidly degraded by the ubiquitin-proteasome system upon with a commercially available HaloPROTAC3 degrader molecule. We demonstrate that inducible degradation of SOX10-HaloTag faithfully recapitulates the known SOX10 transcriptional program, while also enabling discovery of *NAV2* and *NCKAP5* as novel SOX10 target genes. This study demonstrates that new biological insights into the function of SOX10 can be enabled by targeted degradation.

## Introduction

Melanoma, the deadliest skin cancer, is the fourth and fifth most common cancer in men and women in the United States, respectively^1^. Since the introduction of targeted therapies, such as BRAF and MEK inhibitors, as well as checkpoint inhibitor therapy, the five-year survival rates for metastatic melanoma have dramatically improved. However, treatment efficacy remains limited for a substantial percentage of patients due to the development of resistance and adverse responses to therapy. It therefore remains critical to further elucidate the molecular and genetic alterations that drive melanoma in order to identify novel therapeutic opportunities.

Sex determining region Y-box transcription factor 10 (SOX10) is a lineage-specific transcription factor expressed in melanocytes. SOX10 is often upregulated in melanoma cells, which require it for proliferation^2^, and is thus used as a clinical biomarker^3^. SOX10 also controls immune checkpoint expression^4^ via IRF4-IRF1^5^, and its loss has been reported to sensitize melanoma to killing by cytotoxic immune cells^6,7^. An investigational covalent SOX10 inhibitor that acts as a molecular glue to stabilize a reportedly inactive SOX10 homodimeric complex has recently been reported^8^; however, the biological significance of induced SOX10 dimerization remains unclear. Furthermore, the therapeutic value of SOX10 inhibition or depletion has come into question, because low SOX10 expression does not lead to melanoma cell death, but rather induces a transition into a non-proliferative, mesenchymal, invasive cell state, which promotes drug resistance and distant metastases^9,10^.

The molecular function of SOX10 remains incompletely understood, especially in primary melanoma cells, which more faithfully replicate tumor cells than high-passage, immortalized melanoma cell lines. Furthermore, regardless of the cell line used for experiments, SOX10 is challenging to study with standard molecular and genetic tools. Human SOX10 is predominantly intrinsically disordered outside its DNA-binding high mobility group domain^11^ and thus its structure and detailed molecular mechanism have not been elucidated. Furthermore, because SOX10 silencing or deletion abolishes melanoma cell proliferation^9^, it is difficult to separate gene expression changes directly resulting from SOX10 depletion from secondary effects of dormancy. These challenges continue to limit our understanding of the transcriptional program controlled by SOX10 and the regulatory roles it plays in the hallmark pathways of melanoma progression. Importantly, new technologies show promise in addressing some of these gaps in our understanding of SOX10 function and mechanism. For example, a recent study^12^ utilized proximity proteomics to generate the first comprehensive map of the protein communities associated, both directly and indirectly, with SOX10 in melanoma cells.

In this study, we developed a strategy for inducible degradation of SOX10 in primary melanoma cells. Because no proteolysis targeting chimera (PROTAC) for endogenous SOX10 has been reported to date, we leveraged HaloTag-mediated proteolysis targeting chimeras (HaloPROTACs)^13^. HaloPROTACs are bifunctional molecules that bear a covalent ligand for the HaloTag protein linked to an E3 ubiquitin ligase ligand. HaloTag fusion proteins can be readily degraded by the proteasome when the HaloPROTAC, which facilitates the polyubiquitination of the HaloTag fusion protein by the E3 ligase through induced proximity, is added to cells (Figure 1A). HaloPROTACs bear features and mechanism in common with PROTACs under clinical development, and have previously been applied to proof of concept or functional studies of G3BP1^14^ and SCRIB^15^. This approach avoids the drawbacks of SOX10 silencing or deletion, which induce dormancy and may result in off-target effects, and has the advantage over the recently reported SOX10 molecular glue of eliminating SOX10 protein instead of sequestering it in a homodimeric complex. We utilized HaloPROTAC-mediated degradation, coupled with validation using multiple orthogonal datasets, to define the SOX10 transcriptional program in early-passage, patient-derived melanoma cells. This approach enabled the identification and validation of several novel SOX10 target genes, which we show do not reach statistical significance when SOX10 is depleted with siRNA. This study provides new insights into the genes controlled by SOX10 in melanoma cells, while also developing a cell line and approach that can be more broadly applied to the study of SOX10 and other transcription factors.

**Figure 1:**
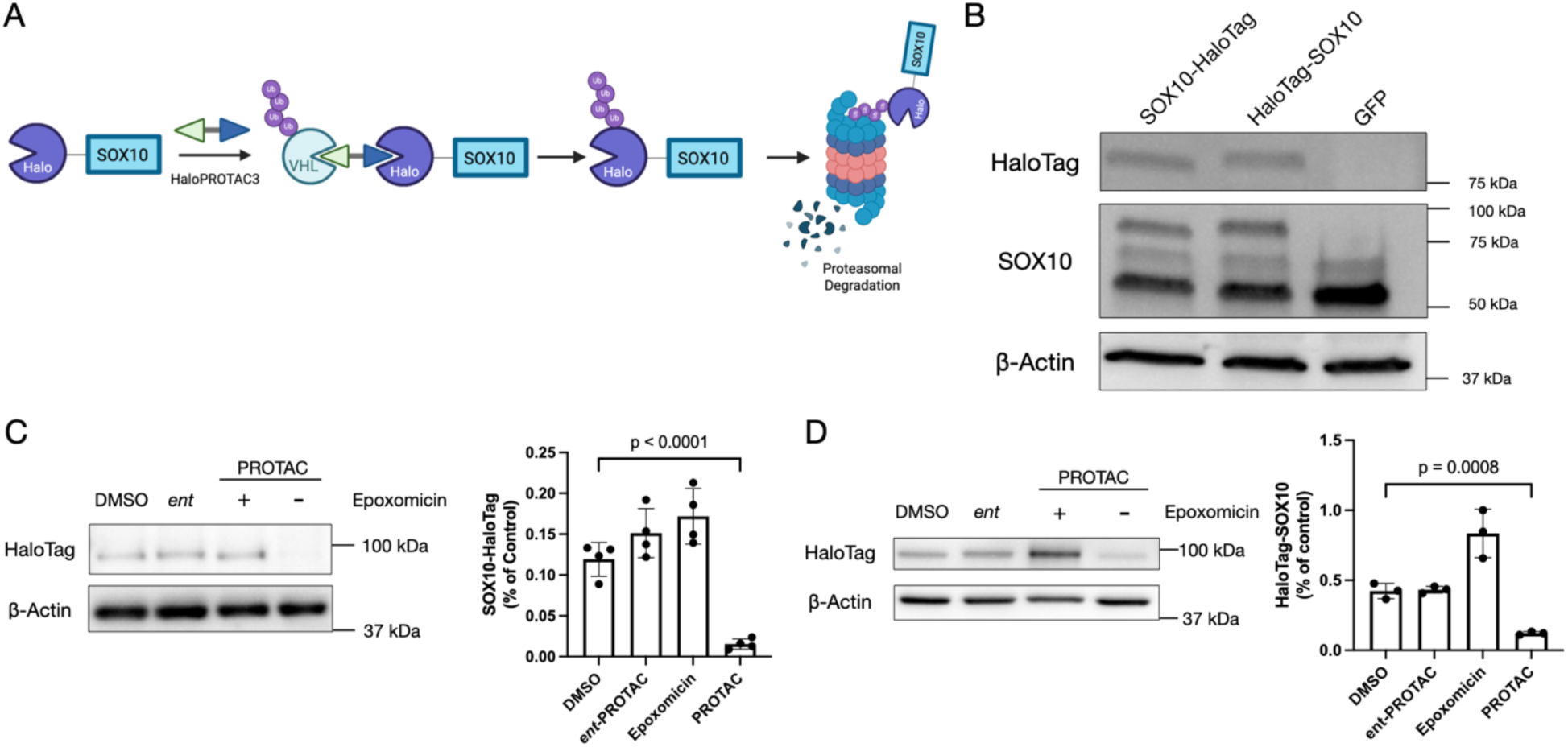
HaloPROTAC-mediated SOX10 fusion protein degradation in melanoma cells. (A) Schematic of SOX10-HaloTag (Halo) fusion protein degradation in the presence of HaloPROTAC3 by the ubiquitin (Ub)-proteasome system. (B) Stable SOX10 fusion protein expression in WW165 melanoma cells assessed with Western blotting. HaloTag-SOX10, N-terminal HaloTag fusion to SOX10; SOX10-HaloTag, C-terminal fusion of HaloTag to SOX10. A plasmid encoding GFP in place of SOX10-HaloTag was stably introduced in WW165 as a negative control (GFP). (C/D) Validation of proteasome-mediated targeted degradation by HaloPROTAC3 (PROTAC) of the SOX10-HaloTag (C) and HaloTag-SOX10 (D) fusion constructs using Western blotting (left). *ent*, diasteromeric *ent*-HaloPROTAC3 (*ent*-PROTAC). Band densitometry of SOX10 fusion protein levels (right). Bars represent the mean value of 4 biological replicates for SOX10-HaloTag and 3 biological replicates for HaloTag-SOX10. Error bars represent standard deviation. P-values, unpaired, two-tailed t-test. β-actin, loading control. Figure prepared in part using BioRender.

## Results

### Engineering patient-derived melanoma cells to express a SOX10-HaloTag fusion protein for inducible degradation

We sought to design a system for on-demand SOX10 degradation in melanoma cells. We selected patient-derived WW165 cells, which are early-passage cutaneous melanoma cells bearing an activating BRAF^V600K^ mutation^16^. Our first goal was to replace the genomic copies of SOX10 with a SOX10 transgene fused to HaloTag. We reasoned that the HaloTag-SOX10 fusion protein would support cell growth and proliferation upon deletion of endogenous SOX10, while enabling conditional SOX10 degradation with enhanced temporal control when cells are treated with HaloPROTAC (Figure 1A).

We first tested expression and degradation of SOX10-HaloTag fusion proteins in wild-type WW165 cells. The SOX10 N-and C-termini present different environments^12^, so we prepared fusions of HaloTag to both termini of SOX10 with an intervening flexible linker. Both constructs expressed at similar levels when stably introduced into WW165 cells (Figure 1B) and were readily degraded by addition of a previously reported, optimized HaloPROTAC compound, HaloPROTAC3^13^ (Figures 1C-D). The presence of HaloPROTAC3, E3 ligase, and the proteasome were all required for SOX10 degradation, as expected. A diastereomeric PROTAC analog that does not bind to the E3 ligase did not support degradation of either SOX10 fusion protein and the addition of a proteasome inhibitor prevented both SOX10 fusion proteins from degradation by HaloPROTAC3, thereby confirming degradation was mediated by the ubiquitin-proteasome pathway. Both fusion constructs could be degraded by HaloPROTAC3 within 24 hours at an optimal concentration of 1.25 µM, consistent with prior reports^13^ (Figure S1). We concluded that both N- and C-terminal SOX10 fusions to HaloTag efficiently undergo induced degradation.

We next deleted the genomic copies of SOX10 from WW165 cells stably expressing the N- or C-terminal fusions of SOX10 to HaloTag. We utilized a doxycycline-inducible Cas9 expression cassette^17^ to minimize off-target genome editing. To prevent Cas9 cleavage of the stably expressed SOX10-Halotag transgenes, the construct underwent codon optimization to avoid nucleic acid sequence complementarity to the guide RNAs without altering the amino acid sequence of encoded SOX10 (Figure 2A). After doxycycline induction, single cell sorting and colony genotyping, we successfully isolated a monoclonal WW165 cell line expressing the C-terminal fusion construct (hereafter, SOX10^HaloTag/KO^) in which all genomic copies of SOX10 were deleted (Figure 2B). The SOX10^HaloTag/KO^ cell line grew in culture despite exhibiting slightly decreased proliferation relative to parental WW165 cells (Figure S2A). SOX10^HaloTag/KO^ cells over-expressed the SOX10-HaloTag transgene by a factor of ∼7-fold compared to endogenous SOX10 levels in wild-type WW165 cells, likely to compensate for loss of endogenous SOX10 (Figure S2B). Stable expression of SOX10-HaloTag is thus functional and supports WW165 cell proliferation in the absence of endogenous SOX10.

**Figure 2:**
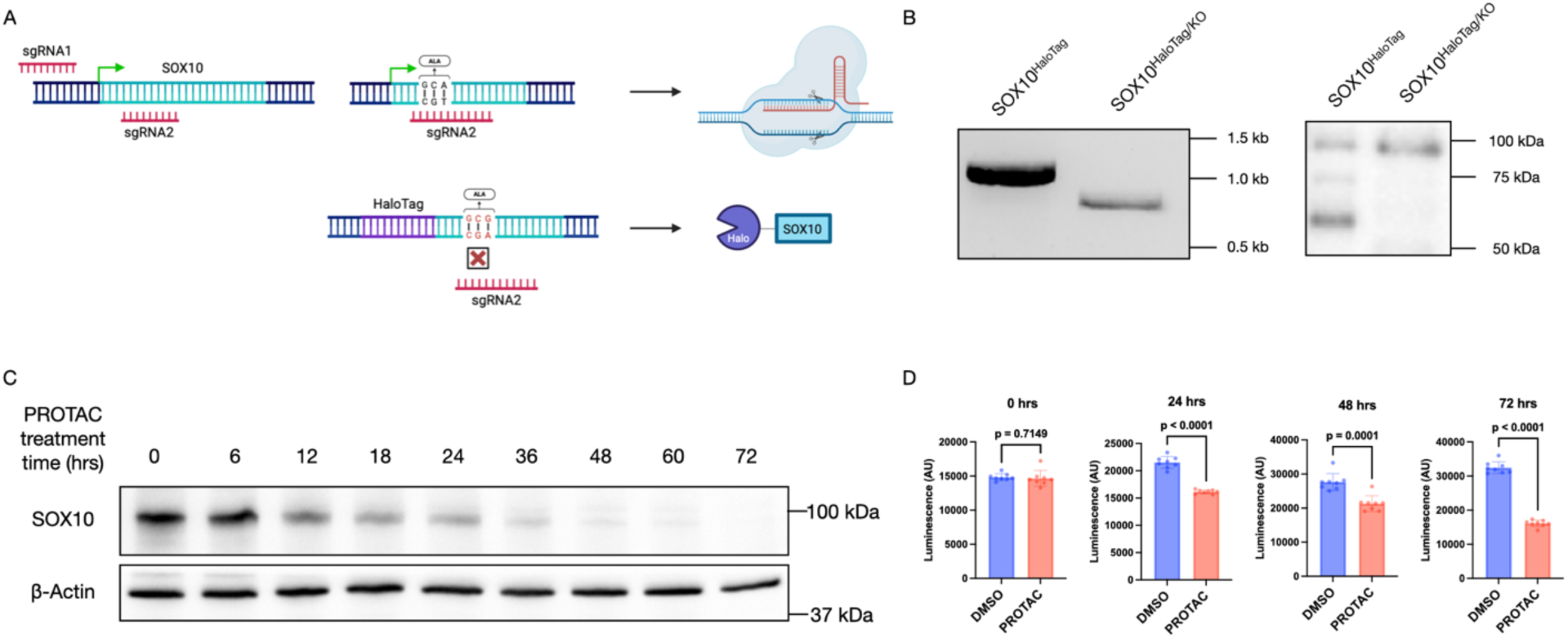
Generation of SOX10 knockout (KO) WW165 cells expressing SOX10-HaloTag (SOX10^HaloTag/KO^). (A) Schematic of CRISPR editing strategy. Single guide RNAs (sgRNAs) that recognize the genomic sequence of SOX10, but not the stably expressed, codon-optimized SOX10-HaloTag construct, were used for SOX10 KO. (B) Validation of endogenous SOX10 KO, and retention of SOX10-HaloTag expression, in SOX10^HaloTag/KO^ cells with genomic PCR (left) and SOX10 Western blot (right). (C) Time course of HaloPROTAC3-mediated degradation of SOX10-HaloTag in SOX10^HaloTag/KO^ cells assessed via SOX10 Western blot. β-actin, loading control. (D) Cell proliferation assay in SOX10^HaloTag/KO^ cells treated with DMSO, blue, or HaloPROTAC3 (PROTAC), red. Error bars, standard deviation (n=8); p-values, unpaired, two-tailed t-test. Figure prepared in part using BioRender.

Finally, we confirmed that SOX10-HaloTag degradation by HaloPROTAC3 proceeds efficiently in SOX10^HaloTag/KO^ cells. HaloPROTAC3 treatment led to ∼67% SOX10-HaloTag degradation by 24 hours, ∼95% by 48 hours, and complete degradation of SOX10-HaloTag within detection limits by 72 hours (Figure 2C). As expected, treatment with HaloPROTAC3 inhibited proliferation of SOX10^HaloTag/KO^ but not wild-type WW165 cells (Figure 2D). In summary, we engineered a patient-derived melanoma cell line in which the only functional copy of SOX10 is fused to HaloTag and can be eliminated upon treatment with HaloPROTAC3.

### Identification of SOX10 target genes in WW165 cells

We applied HaloPROTAC3-mediated SOX10 degradation to high-sensitivity mapping of the SOX10 transcriptional program. We first confirmed that HaloPROTAC3 on its own does not artefactually alter gene expression in WW165 cells. We carried out a control experiment in which parental WW165 cells were treated with HaloPROTAC3 or vehicle, followed by RNA-seq. A single gene exhibited a small but statistically significant decrease in the HaloPROTAC3-treated samples; all other genes were unchanged (Figure S3). We therefore concluded that HaloPROTAC3 treatment alone has little effect on the WW165 transcriptome and thus is compatible with SOX10 functional studies.

We then quantified changes in gene expression in SOX10^HaloTag/KO^ cells after SOX10-HaloTag degradation compared to vehicle treatment (Figure 3A, Table S2). We observed statistically significant downregulation of more than 700 genes, including well-validated SOX10 target genes like *MITF*, *ERBB3*, and *DCT*^18,19^ (Figure 3B). The downregulated genes were associated with a number of gene ontology (GO) terms related to cell division, consistent with inhibition of cell proliferation by SOX10 depletion (Figure S4). We next examined changes in chromatin accessibility using the assay for transposase accessible chromatin (ATAC-seq) under the same conditions (Figure 3B, Table S2). Of 3000 genes exhibiting decreased accessibility after SOX10 degradation, 101 were also downregulated in the RNA-seq dataset, again including true positive SOX10 targets. The convergence of decreased transcription and chromatin accessibility strongly support the likelihood that these 101 intersectional genes require SOX10, either directly or indirectly, for their expression in WW165 cells (Figure 3B). We thus consider the set of 101 genes to be stringent candidates for SOX10 regulation; however, there are likely many additional *bona fide* SOX10 target genes in each dataset that did not meet our thresholds. We also note that we did not investigate genes that are upregulated by SOX10 degradation, though these may also be biologically significant. These datasets thus provide a resource for further study of SOX10-dependent transcriptional changes in melanoma.

**Figure 3.**
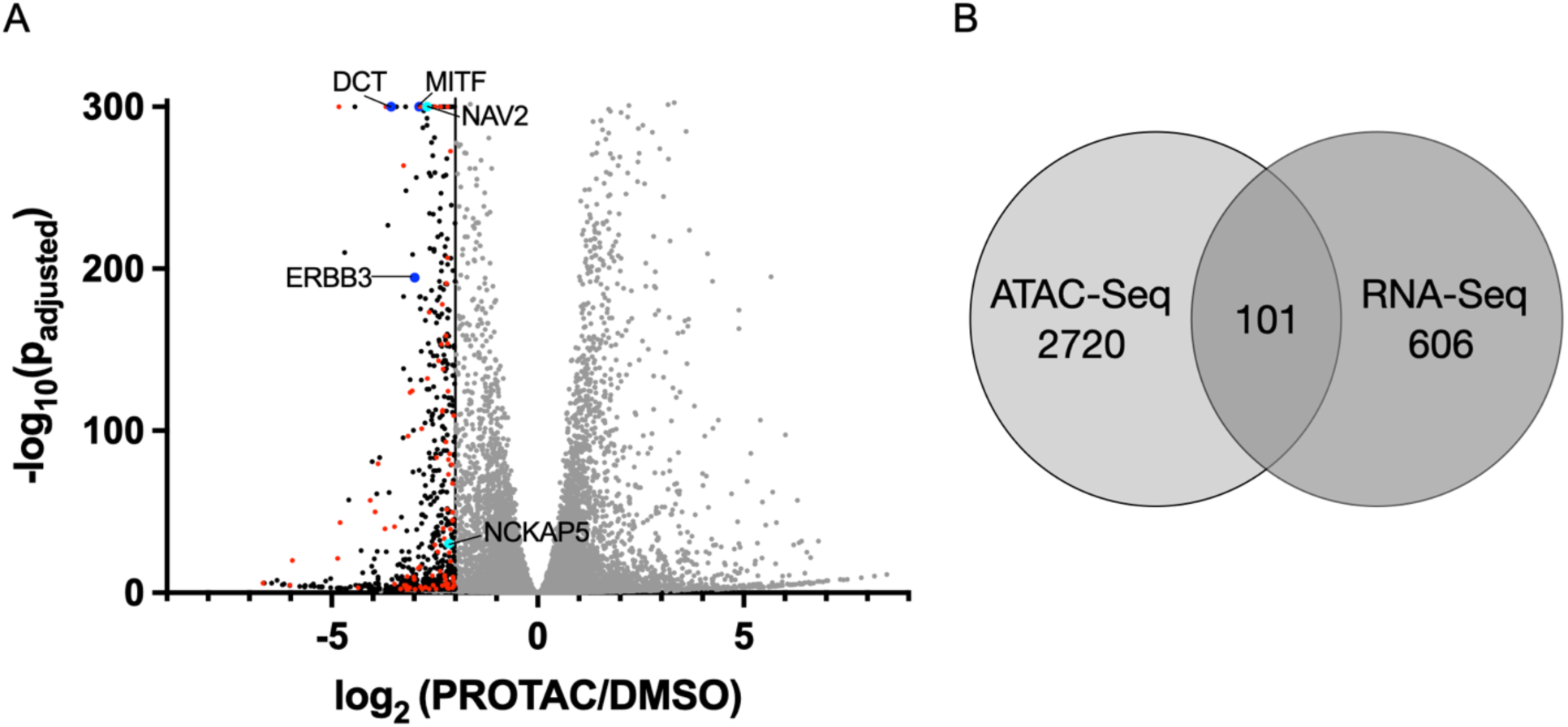
Integrative profiling of transcriptional changes in WW165 melanoma cells after SOX10-HaloTag degradation. (A) Differential expression analysis of RNA-seq data from SOX10^HaloTag/KO^ cells treated with HaloPROTAC3 (PROTAC) or DMSO for 48 hours. Black data points are downregulated genes defined by log_2_(PROTAC/DMSO) ≤ –2 with a p_adjusted_ value ≤0.01. Red data points are genes significantly downregulated after SOX10-HaloTag degradation by both RNA-seq and ATAC-seq. Blue data points are previously reported SOX10 target genes. Cyan data points are validated (vide infra) novel SOX10 target genes. (B) Venn diagram of overlapping and unique genes downregulated by SOX10-HaloTag degradation in SOX10^HaloTag/KO^ cells in both RNA-seq and ATAC-seq analyses. Significant genes were defined as log_2_(PROTAC/DMSO) ≤ –2 with a p_adjusted_ value ≤0.01 for each dataset.

We next compared our method to SOX10 knockdown. In our hands, siRNA transfection afforded ∼ 73% SOX10 silencing in wild-type WW165 cells. We performed RNA-seq after SOX10 knockdown (Figure 4A) and compared differentially expressed genes to those downregulated after HaloPROTAC-mediated degradation. We observed 571 significantly downregulated genes in siRNA-treated cells, and 707 in HaloPROTAC3-treated SOX10^Halo/KO^ cells. 282 genes were significantly downregulated in both datasets (Figure 4B). Included among the shared genes were well-established SOX10 targets, supporting the validity of both experimental approaches for SOX10 depletion. However, each approach also yielded hundreds of differentially expressed genes that were not detected in the other experiment. We hypothesized that some of the genes downregulated specifically by SOX10-HaloTag degradation could represent *bona fide* SOX10 target genes that did not reach the cutoff for effect size or significance after siRNA treatment. Supporting this hypothesis, differentially expressed genes after either siRNA or HaloPROTAC3 treatment exhibited a correlation coefficient of 0.66 (Figure 4C). A majority of the genes that were significantly downregulated only in the HaloPROTAC3 treatment condition were also downregulated after siRNA treatment but failed to reach statistical significance in the latter condition. This suggests that the regulation of some genes can only be confidently detected when SOX10 is degraded; we speculate that this could be a result of more complete or rapid SOX10 protein depletion by targeted degradation under the experimental conditions used.

**Figure 4.**
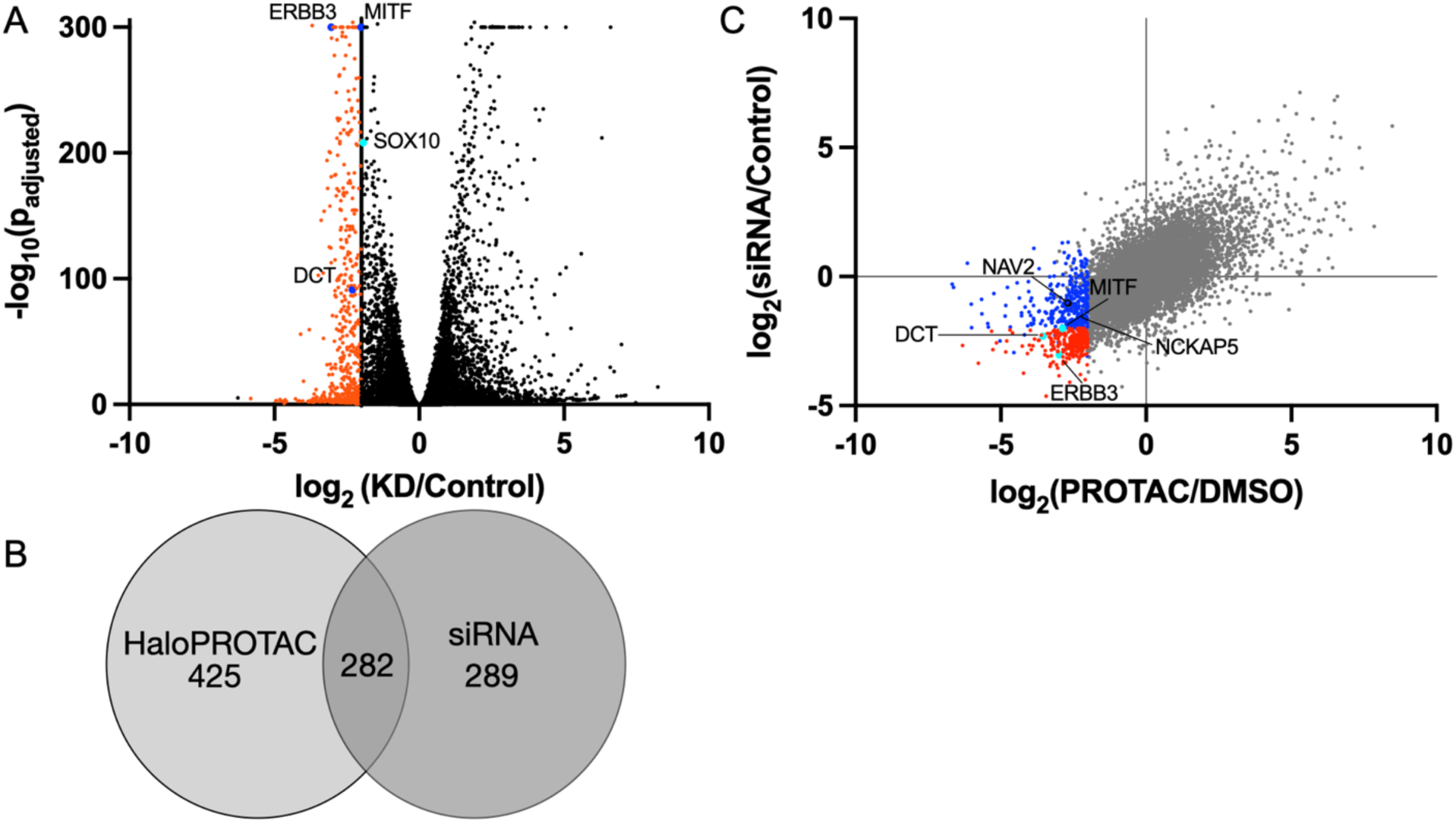
Comparing transcriptional effects of SOX10 silencing to SOX10 degradation. (A) Differential expression analysis of RNA-seq data from wild-type WW165 cells transfected with SOX10 siRNA or negative control siRNA. Red data points are significantly downregulated genes. Blue data points are previously reported SOX10 target genes. Cyan data point is SOX10. (B) Venn diagram of overlapping and unique genes significantly downregulated in SOX10^HaloTag/KO^ cells treated with HaloPROTAC3 (PROTAC) vs. vehicle (from Figure 3A), and in wild-type WW165 cells subjected to SOX10 siRNA vs. control (from Figure 4A). (C) Correlation plot of differential gene expression after SOX10 silencing or HaloPROTAC3-mediated degradation. Red data points represent genes downregulated in both treatments, while blue data points highlight genes significantly downregulated only by HaloPROTAC3-mediated SOX10 degradation. Cyan data points are true positive SOX10 target genes. Black points are validated (vide infra) novel SOX10 target genes. All significantly downregulated genes are defined as log_2_(PROTAC/DMSO) or log_2_(KD/Control) ≤ –2 with a p_adjusted_ value of ≤0.01.

We next determined whether SOX10 degradation enables identification of novel genes that depend on SOX10 for their transcription, either directly or indirectly. We prioritized 13 candidates among the 101 genes downregulated in both the RNA-seq and ATAC-seq experiments after SOX10 degradation that, to our knowledge, have not been previously validated as SOX10 targets. These include *NCKAP5*, *NAV2, TDRD7*, *MSK1*, *CHD1*, *ALK*, *PKNOX2*, *ITGA9*, *CD200*, *RFTN1*, *DKK2*, *MDGA1*, and *ICOS* (Figure 5, Figures S5-S8, Table S2). First, we tested their RNA-level downregulation after SOX10-HaloTag degradation in WW165 cells via RT-qPCR (Figure 5, Figure S5). For 8/13 genes tested, RNA levels decreased with statistical significance after HaloPROTAC3-mediated SOX10 degradation, as expected. Three genes failed to amplify from WW165 cells using previously reported or newly designed primers and two did not exhibit statistically significant decreases (Figure S5, Table S2).

**Figure 5:**
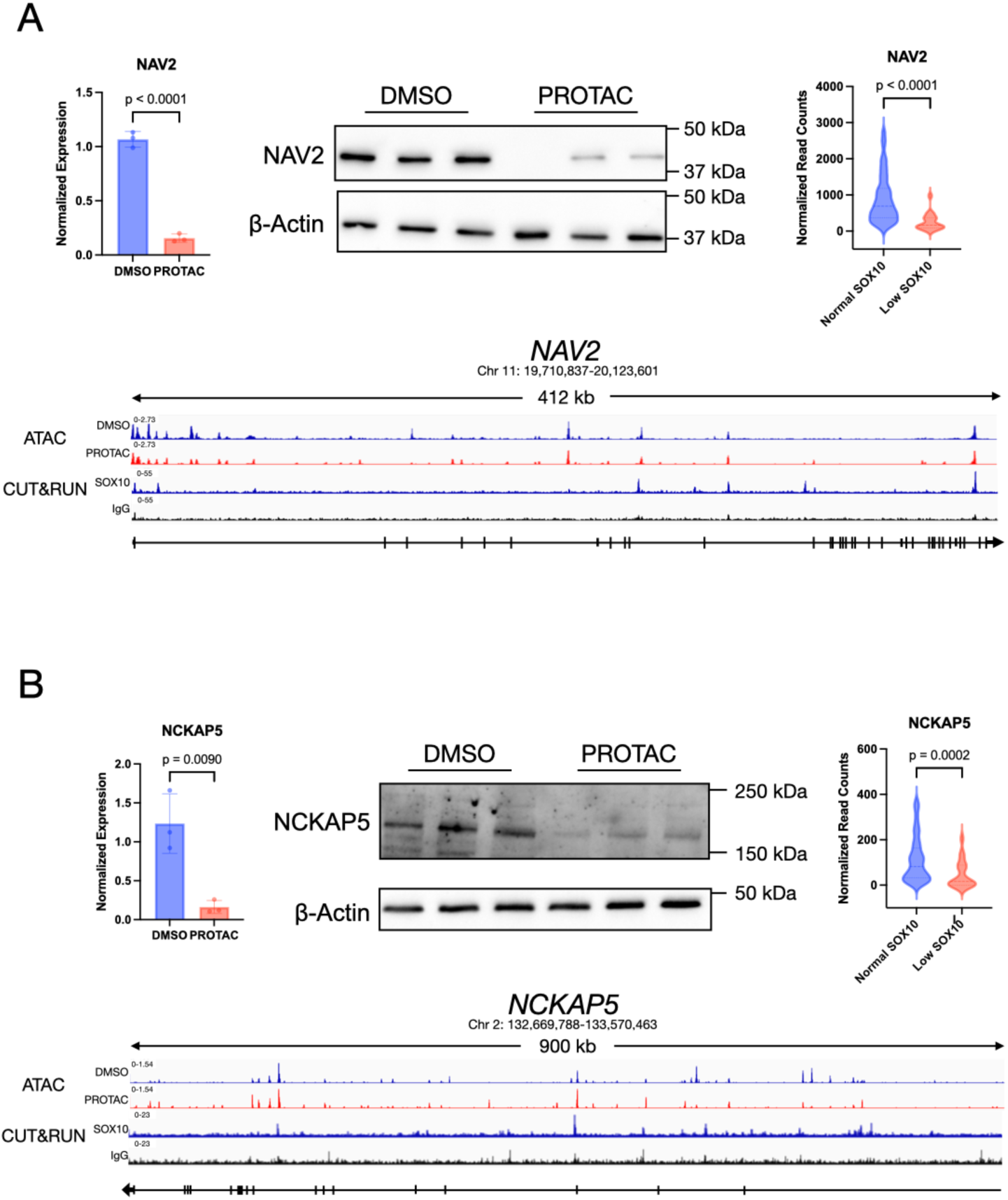
Validation of SOX10 target genes (A) *NAV2* and (B) *NCKAP5*. Top left, RT-qPCR analysis of RNA isolated from SOX10^HaloTag/KO^ cells treated with DMSO or HaloPROTAC3 (PROTAC) for 48 hours. P-values, unpaired t-test; error bars, standard deviation. Each point represents one biological replicate, itself the average of three technical replicates. Top middle, Western bloting of SOX10^HaloTag/KO^ cells treated with DMSO or HaloPROTAC3 (PROTAC) for 48 hours. Each band represents one biological replicate. Top right, normalized read counts for the gene of interest in low-SOX10 expressing patient melanoma cells (normalized SOX10 read counts < 200) to normal-SOX10 expressing patient cells (normalized SOX10 read counts > 200). SOX10 expression data were filtered using the ROUT test to remove outliers followed by a non-parametric Mann-Whitney test for significance (Q=1%). P-values, Mann-Whitney test. Bottom, ATAC-seq and CUT&RUN. Top two tracks, ATAC-seq from SOX10^HaloTag/KO^ cells treated with DMSO (blue) or HaloPROTAC3 (red) for 48 hours. Signal represents the average of 3 biological replicates. Bottom two tracks, reanalysis of SOX10 CUT&RUN signal (blue) vs. IgG control (black)^20^. CUT&RUN signal represents the average of 3 replicates.

We next examined protein-level regulation of novel target genes by SOX10. We obtained specific antibodies to *NAV2*, *NCKAP5, MSK1*, *CD200*, and *RFTN1* and confirmed that that these 5 proteins are downregulated after SOX10-HaloTag degradation (Figure 5, Figure S6, and Table S2). *DKK2* and *PKNOX2* did not exhibit protein-level decreases after SOX10 degradation (Figure S6). We were either unable to procure a specific antibody, or did not observe specific bands on Western blots, corresponding to *TDRD7*, *CHD1*, *ALK*, *ITGA9*, *MDGA1*, or *ICOS*. We thus validated transcriptional downregulation of 9 novel genes after SOX10 degradation and confirmed protein-level downregulation of 5. We conclude that expression of these 9 genes in WW165 cells depends on SOX10, either via direct binding or through secondary signaling pathways, and that this SOX10 dependency controls protein levels for at least 5.

Our system relies on overexpression of a SOX10 transgene; it was therefore essential to determine whether the putative target genes we identified are directly bound by endogenously expressed SOX10. To do so, we re-examined previously reported SOX10 CUT&RUN data from human induced pluripotent stem cell (hPSC)-derived melanocytes^20^. As shown in Figure 5, Figure S7, and Table S2, 10 of the 13 examined target genes exhibit SOX10 CUT&RUN peaks above IgG background at or near the promoter region or within the gene body. By comparison, a positive control gene, *MITF*, exhibited specific SOX10 signal (Figure S7, top left), while a negative control gene, *TNNI3*, which is not expressed in skin, had low signal comparable to IgG background (Figure S7, bottom right). We then examined ATAC-seq tracks from our dataset (Figure 3B) for the same genes after SOX10-HaloTag degradation and found that, in general, decreased chromatin accessibility after SOX10-HaloTag degradation was consistent with SOX10 binding (Figure 5, Figure S7). We conclude that a majority of genes downregulated by SOX10 degradation in our system may be bound by endogenous SOX10 in hPSC-derived melanocytes, consistent with transcriptional regulation.

Lastly, we examined whether the 13 candidate target genes are regulated by SOX10 beyond WW165 cells. To do so, we stratified RNA-seq data previously acquired^16^ from hundreds of sun-exposed and sun-shielded melanomas according to SOX10 expression levels (275 cell lines with normal SOX10 expression, 22 cell lines with low SOX10 levels defined as normalized read counts <200, after exclusion of outliers using ROUT test with Q=1%), and examined expression of each of the target genes in low vs. high SOX10 expressing samples (Figure 5, Figure S8, and Table S2). In seven cases (*NAV2*, *NCKAP5*, *ALK*, *TDRD7*, *PKNOX2, CHD1*, *CD200*), low SOX10 expression significantly correlated with decreased target gene levels, suggesting that SOX10 is associated with transcription of these genes in melanoma. Five genes were not decreased in low SOX10-expressing samples, and one was not detected (Figure S8, Table S2).

Integrating our RNA-seq and ATAC-seq results after SOX10-HaloTag degradation with RT-qPCR, Western blotting, SOX10 CUT&RUN, and patient expression data, we concluded that *NKCAP5* and *NAV2* were fully validated as SOX10 target genes (Figure 5). *TDRD7* lacked specific antibodies for Western blotting but was supported as a SOX10 target gene by all remaining levels of evidence. The other 10 novel genes examined were associated with SOX10 at multiple levels but lacked at least one level of validation, and thus may represent either direct or indirect targets of SOX10. Our method can thus reveal novel, cancer-relevant SOX10 genes and effectively map a portion of the transcriptional program previously missed by traditional experimental methods.

## Discussion

In this work, we engineered a patient-derived melanoma cell line to express a SOX10-HaloTag fusion protein that can be degraded upon treatment with a commercially available HaloPROTAC. The SOX10 fusion to HaloTag at either the N- or C-terminus enables HaloPROTAC3-inducible degradation of the fusion protein via the ubiquitin-proteasome pathway within 24-72 hours. The C-terminal fusion protein supports melanoma cell growth when the genomic copies of *SOX10* are deleted, demonstrating that the SOX10-HaloTag fusion protein is functional. This system is complementary to genetic approaches to SOX10 silencing or deletion. The first advantage to our method is that SOX10 degradation is conditional on the addition of the HaloPROTAC, avoiding the disadvantages of constitutive SOX10 ablation; second, HaloPROTAC treatment leads to essentially complete SOX10 degradation within detection limits within the same time frame over which siRNA leads to only partial SOX10 depletion.

We applied HaloPROTAC-mediated SOX10 degradation to high-sensitivity profiling of the SOX10 transcriptional program in patient-derived melanoma cells. Targeted degradation of SOX10-HaloTag followed by multidimensional profiling accurately identified well-known SOX10 target genes (e.g., *MITF*, *ERBB3*, and *DCT*) and processes (e.g., cell cycle regulation), demonstrating that this method identifies *bona fide* SOX10 targets. Intersection of our RNA-seq and ATAC-seq datasets prioritized 101 high-confidence genes that require SOX10 for their transcription in WW165 cells. These SOX10-regulated genes are associated with cell cycle progression, consistent with the known essential role of SOX10 in melanoma cell proliferation. Importantly, many of the SOX10 target genes identified by our method are also downregulated when cells are treated with siRNA targeting SOX10, but do not reach statistical significance in the latter condition, suggesting that targeted degradation of SOX10 affords enhanced sensitivity for detection of a subset of SOX10-regulated genes.

We identified a number of novel SOX10 target genes using this methodology. Importantly, we validated *NAV2* and *NCKAP5* as direct SOX10 target genes via a series of downstream analyses, demonstrating that they are not only downregulated by SOX10 targeted degradation, but are also bound by endogenous SOX10 and are coregulated with *SOX10* levels in a patient cohort. These genes have previously been linked to melanoma initiation, progression, or outcomes, but to our knowledge have not been validated as SOX10 targets. *NAV2*, or neuron navigator 2, encodes a protein involved in cell migration that has been linked to poor prognosis in cutaneous melanoma and promotes increased melanoma cell proliferation and invasiveness *in vitro*^21^. *NCKAP5*, or NCK associated protein 5, was recently implicated in the genetic mechanism by which exposure to UVB radiation overcomes oncogene-induced senescence associated with the activating *BRAF^V600E^*mutation^22^. In the same study, silencing of *NCKAP5*, along with several other genes in the same UV radiation-induced transcriptional reprogramming signature, reduced melanoma cell proliferation. Our demonstration that *NAV2* and *NCKAP5* directly depend on SOX10 for their transcription thus provides a mechanism for their regulation in melanoma cells, and is consistent with known roles for SOX10 in promoting melanoma progression and invasiveness^23^.

While we utilized HaloPROTAC degradation of a SOX10 fusion protein to investigate its transcriptional program in patient-derived melanoma cells, we propose that this system has applications beyond the current study. The HaloPROTAC methodology could be employed to identify targets of other transcription factors – or, more broadly, functions of other essential proteins – with enhanced sensitivity. Furthermore, we imagine that the WW165^KO/HaloTag^ cells generated in this study could be applied to investigate the consequences of SOX10 degradation on immune signaling, HLA I display of epitopes derived from SOX10 after targeted proteolysis, or tumor inhibition *in vivo*. However, our study also has limitations. The SOX10-HaloTag transgene only partially rescues cell proliferation when endogenous SOX10 is ablated, suggesting that it does not fully recapitulate the function of wild-type SOX10. Future work could address this limitation by optimizing the fusion protein, linker, and expression level of the SOX10-HaloTag transgene, or utilizing an alternative degron system^24–26^. Alternatively, conversion of a recently reported SOX10 covalent ligand^8^ into a PROTAC could enable targeted degradation of endogenous SOX10, circumventing the use of fusion proteins altogether.

In conclusion, we have demonstrated HaloPROTAC-mediated degradation of a SOX10 transgene in low-passage melanoma cells. Given sustained interest in SOX10 as a potential therapeutic target in melanoma, this system may be of interest to the broader scientific community for further proof-of-principle studies of the outcomes of SOX10 targeted degradation *in vitro* and, potentially, *in vivo*.

## Acknowledgment

This work was supported in part by a Falk Medical Research Trust Transformational Award (S.A.S. and R.H.), a Melanoma Research Alliance (MRA) Established Investigator Award (R.H. and S.A.S), a MRA Next Steps: Biomarkers Grant (S.A.S.), the Mark Foundation for Cancer Research Emerging Leader Award (to S.A.S.), a Sloan Research Fellowship FG-2022-18417 (S.A.S.), the Yale SPORE in Skin Cancer Developmental Research Project Award (S.A.S.), NIH grant P50CA121974 (the Yale SPORE in Skin Cancer (R.H.), and the Roslyn and Jerome Meyer Award (R.H.). A.L.P., C.M.D. and K.J. were supported in part by NIH grant 1T32GM149444-01. C.M.D. was supported in part by a Gruber Science Fellowship from the Gruber Foundation. All sequencing was performed by the Yale Center for Genome Analysis (YCGA), supported by the National Institute of General Medical Sciences of the National Institutes of Health under Award Number 1S10OD030363-01A1.

## Methods

### Cell Culture

All melanoma cell lines used in this study were cultured as previously reported^27^ in Opti-MEM supplemented with 5% Fetal Bovine Serum (FBS), 1% Penicillin-Streptomycin (PS), and 0.1 mM IBMX (3-isobutyl-1-methyl xanthine, Sigma, I-5879) in a humidified incubator at 37°C and 5% CO_2._ Cells were maintained by consistent splitting once 95% confluency was reached and frequent media changes every 2-3 days. All cell stocks were stored in 90% FBS and 10% DMSO in liquid nitrogen. Upon revival, cells were passaged at least twice prior to any experiments. HEK293T cells were cultured in DMEM supplemented with 10% FBS and 1% penicillin-streptomycin in a humidified incubator at 37°C and 5% CO_2._ All cell lines underwent regular mycoplasma testing using the InvivoGen MycoStrip® test and the manufacturer’s protocol.

### Lentivirus Generation and Transduction

The lentiviral packaging plasmids (9 µg ViraPower, ThermoFisher, K497500) and pLJM1 HaloTag-SOX10 (3 µg) were combined in 1.5 mL of OptiMEM before incubating at room temperature for 5 minutes. Lipofectamine 2000 (36 µL, ThermoFisher, 11668027) was mixed with 1.5 mL OptiMEM prior to mixing with the DNA and then incubating at room temperature for 20 minutes. After incubation, the lipid DNA complex was added to 6 million HEK293T cells resuspended in OptiMEM with 10% FBS and then plated in a 10 cm dish. After 24 hours, the media was replaced with 7 mL OptiMEM with 10% FBS and grown for 24 hours. Lentiviruses were collected 48 hrs and 72 hrs post transfection and stored on ice at 4°C prior to consolidation. After the final collection, any remaining cells were pelleted and the media was passed through a sterile 0.45 µm filter into a 100 kDa spin filter. The viral particles were concentrated by centrifuging for 2,500 RPM for 30 minutes at 10°C. The concentrated virus was then used immediately for transduction or stored at -80°C for future use.

For viral transduction, WW165 cells were grown to 80% confluency in a 10 cm dish. The expired media was aspirated before being replaced with OptiMEM. Polybrene (final concentration of 5 µg/mL) was mixed with 50 µL of lentivirus prior to dropwise addition to the cells. After 24 hours, the media was replaced with OptiMEM supplemented with 5% FBS, 1% PS, and 0.1 mM IBMX and allowed to grow for an additional 24 hours. At 48 hours post transduction, cells underwent puromycin selection at a concentration of 2.5 µg/mL for 72 hours before slowly tapering the puromycin.

### HaloPROTAC3 Treatment

WW165 cells overexpressing HaloTag-SOX10 or SOX10-HaloTag were plated in 12-well plates at a density of 2.5 x 10^5^ cells/mL and allowed to grow for a minimum of 24 hours to a confluency of 70-80%. For HaloPROTAC3 concentration optimization, media was aspirated and replaced with HaloPROTAC3 (Promega, GA3110) supplemented growth media at concentrations ranging from 5 µM to 1.22 nM. An equal volume of DMSO was added to growth media as vehicle control. Cells were incubated for 24 hours at 37°C before being collected and washed with PBS+PAO (phenylarsine oxide). For treatment time optimization,1.25 µM HaloPROTAC3 or DMSO was supplied in fresh growth media at 37°C ranging from 1 hour to 24 hours prior to collection with PBS+PAO. To confirm that degradation of SOX10 fusion proteins requires E3 ligase and proteasome activity, cells were treated with media supplemented with either DMSO as a negative control, 1.25 µM *ent*-HaloPROTAC3 (Promega, GA4110), 1.25 µM HaloPROTAC3 with 0.3 µM epoxomicin, or 1.25 µM HaloPROTAC3 as a positive control, followed by a 24-hour incubation period, washing with PBS+PAO and collection.

For all experiments, cells were lysed by resuspending in SDS protein loading buffer (10% glycerol, 60 mM Tris-HCl pH 6.8, 0.8 g SDS, 1.25% 2-mercaptoethanol, and 4 mg bromophenol blue) and boiling for 5 minutes. HaloTag-SOX10 degradation was assessed by Western blotting using a HaloTag® Monoclonal antibody (Promega, G9211) and beta-Actin Loading Control Monoclonal antibody (Invitrogen, BA3R).

### Western Blotting

All samples were prepared in SDS protein loading buffer followed by boiling at 95°C. Samples were loaded on a 10% polyacrylamide glycine gel and ran at 120 V for approximately 45-60 minutes or until the dye front was 0.5 inches from the bottom of the gel. Proteins were then transferred to a 0.2 µm nitrocellulose membrane over 1 hour at 300mA. The membrane was washed with TBST (Tris Buffered Saline with Tween-20) prior to blocking with 3% BSA (Bovine Serum Albumin) in TBST at room temperature while gently rocking. After 1 hour, the blot was washed with TBST three times prior to primary antibody incubation overnight at 4°C. Primary antibodies were made 1:1,000 in 3% BSA in TBST unless stated otherwise. After washing with TBST three times, the blot was incubated for 1 hour at room temperature with an HRP-conjugated secondary antibody at 1:10,000 in 3% BSA in TBST. The blot was washed with TBST, a 1:1 solution of hydrogen peroxide and luminol (BioRad Clarity Western ECL Substrate, 1705061) was added and incubated at room temperature for 5 minutes before imaging.

### CRISPR/Cas9 SOX10 Knockout in WW165 Cells

CRISPOR (UC Santa Cruz, Version 5.2) was used to generate single guide RNA (sgRNA) sequences targeting the start codon of endogenous SOX10, which were cloned into TLCV2 (Addgene plasmid #87360) as previously reported. sgRNA sequences are provided in Table S1. Lentiviruses expressing each sgRNA sequence were generated in HEK 293T as described above.

WW165 cells stably expression HaloTag-SOX10 or SOX10-HaloTag were seeded in a 6-well plate and allowed to grow for at least 24 hours prior to transduction. Once cells reached 70-80% confluency, expired media was aspirated prior to being replaced with Opti-MEM supplemented with 5 µg/mL polybrene. A total of 80 µL of 1:1 forward sgRNA and reverse sgRNA lentiviruses were added dropwise to the cells and allowed to incubate for 24 hours prior to induction. Expired viral media was removed and replaced with Opti-MEM supplemented with 0.75 µg/mL doxycycline and incubated at 37°C for 24 hours.

Following Cas9 induction with doxycycline, cells were detached from the plate with trypsin before being collected and resuspended in 1 mL of FACS growth media (Opti-MEM, GlutaMAX, 5% FBS, 1% PS, and 0.1 mM IBMX). Cells were then passed through a cell strainer before being sorted for EGFP expression in 96-well plates at a density of 1 cell/well. FACS was performed using a ThermoFisher BigFoot spectral cell sorter with transduced cells in the absence of doxycycline induction as a negative control. Cells were then allowed to grow at 37°C for approximately 1 month or until at least 80% confluency.

Once cells had reached at least 80% confluency, the cells were split by half into a new 96-well plate to be used for monoclonal cell line propagation. Remaining cells were lysed by removing the growth media and adding 50 µL of 10 mM Tris/HCl pH 8, 0.05% SDS, and Proteinase K, followed by genomic DNA isolation and PCR using the primer sequences shown in Table S1. To further validate SOX10 knockout, TOPO cloning was performed using the TOPO TA Cloning Kit (Invitrogen, 450071) following the manufacturer’s protocol and subjected to Sanger sequencing.

### Cell Proliferation Assay

Cells were plated at a density of 4,000 cells/well in white bottom 96-well plate and incubated at 37°C. At each timepoint, plates were removed from the incubator and media was removed. To each well, 50 µL of reconstituted Cell Titer Glo (Promega, G7570) was added to each well and mixed well by pipetting. The plate was covered with aluminum foil followed by gently rocking the plate for 2 minutes at room temperature. The plate was removed from the rocker and allowed to incubate for 10 minutes at room temperature prior to imaging. Luminescence values were normalized to a background well containing media and reconstituted Cell Titer Glo.

### RNA-seq and Data Analysis

For analysis of transcription changes after SOX10 degradation, SOX10^HaloTag/KO^ cells at ∼80% confluency were treated with 1.25 µM of HaloPROTAC3 or an equal volume of DMSO for 48 hours in standard melanoma cell culture media, followed by washing with DPBS. Cells were collected and processed for RNA extraction as below.

For analysis of transcription changes after SOX10 knockdown, wildtype WW165 cells were cultured in T-25 flasks in standard culture media until 70% confluency was achieved. Cells were transfected with SOX10 targeting siRNAs (Ambion s13309/s13310), or *Silencer*™ Select Negative Control No. 1 siRNA (Invitrogen, 4390843) as a negative control, at a final concentration of 18.75 nM using Opti-MEM and Lipofectamine RNAiMAX. Cells were incubated for 48 hours prior to washing with DPBS and collection.

After treatment, cells were collected by trypsinization and mechanical detachment and pelleted by centrifugation at 1,300 RPM at 4°C. Cells were washed once with PBS and pelleted once more before PBS was removed. RNA was purified using TRIzol (ThermoFisher,15596026) following the manufacturer’s protocol, followed by RNA clean-up using the QIAgen RNeasy Kit (catalog # 74104).

For RNA-seq analysis of each sample type, 1 μg of purified total RNA per replicate was submitted to the Yale Center for Genome Analysis, where ribosomal RNA depletion and library preparation was performed. Sequencing was performed with the Illumina NovaSeq system at a depth of approximately 35 million reads per sample. Paired-end reads were sequenced with a read length of 150 base pairs.

Raw sequencing reads were assessed for quality using FastQC (v0.12.1) and trimmed to remove adapter sequences and low-quality bases using cutadapt (v5.2). Filtered reads were aligned to the human reference genome (hg38) using STAR (v2.7.11b) with permissive multimapping parameters (--twopassMode Basic, --sjdbOverhang 100, -- winAnchorMultimapNmax 200, --outFilterMultimapNMax 100). Alignment files were converted to sorted and indexed BAM format using SAMtools (v1.22.1). Gene-level read quantification was performed using featureCounts (v2.1.1) with exon-level summarization and assignment by gene ID using the GENCODE hg38 gene annotation. Paired-end fragments were counted, and multimapping or ambiguously assigned reads were excluded from downstream analysis. Quality control metrics across all processing steps were aggregated using MultiQC (v1.28).

Downstream analysis was performed in R (v4.4.1). Raw count matrices were filtered to remove lowly expressed genes, retaining genes expressed in at least 50% of samples and with a minimum total count threshold of 24. Differential expression analysis was conducted using the DESeq2 package (v1.44.0), which models count data using a negative binomial distribution and incorporates normalization for sequencing depth and dispersion across biological replicates. P-values were derived from Wald tests and corrected for multiple testing using the Benjamini– Hochberg method. Genes with an adjusted p-value < 0.01 and an absolute log2 fold change > 2 were considered significantly differentially expressed and were used in all subsequent analyses.

### Reverse Transcription and Quantitative Polymerase Chain Reaction (RT-qPCR)

The BioRad iScript cDNA Synthesis Kit was used to reverse transcribe purified RNA for use in qPCR. Each reaction proceeded with 1 µg of total RNA following the manufacturer’s protocol for reaction composition, thermocycler timing, and temperature requirements. For qPCR, Luna Universal qPCR Master Mix (NEB, M3003) was used with forward and reverse primers for the target gene of interest (0.5 µM) and 40 ng of cDNA for a total volume of 10 µL. Primer sequences are provided in Table S1. Thermocycling conditions followed the manufacturers protocol with scans taken using the SYBR/FAM mode on a BioRad CFX96 Real-Time System (C1000 Touch™ Thermal Cycler). Results were processed in BioRad CFX Manager™ Software (Version 3.1) and raw Cq values were normalized to actin before calculating 2^-ΔΔCq^ and performing an unpaired t-test in GraphPad Prism.

### ATAC Sequencing

SOX10^HaloTag/KO^ cells were treated for 48 hours with either 1.25 µM HaloPROTAC3 or DMSO as a vehicle control. The Vazyme Hyperactive ATAC-Seq Library Prep Kit for Illumina (catalog # TD711) was used to tagment, isolate, and amplify the DNA library. Results presented in this paper were generated using 100,000 cells per reaction and the Vazyme ATAC-Seq protocol (Version 24.1) and sequenced at the Yale Center for Genome Analysis.

Raw ATAC-seq reads were processed using the nf-core/atacseq pipeline (v2.1.2) implemented in Nextflow using the human GRCh38 (hg38) reference genome. Briefly, adapter trimming was performed using Cutadapt, reads were aligned using BWA. PCR duplicates and mitochondrial reads were removed during preprocessing. Accessible chromatin regions were identified using MACS2 with narrow peak calling enabled. Sequencing reads were analyzed with a read length parameter of 100bp, and consensus peak sets were generated requiring a minimum of two replicates per condition.

Differential chromatin accessibility analysis was performed in R using the DiffBind package. Differential analysis was performed without additional blacklist or greylist filtering, and all peaks were retained for downstream analysis prior to downstream significance filtering. Differentially accessible peaks were annotated using the ChIPseeker package and the hg38 transcript annotation, with promoter regions defined as ±1 kb from annotated transcription start sites (TSSs). For downstream analyses, differentially accessible ATAC-seq peaks were filtered using thresholds of false discovery rate (FDR) < 0.01 and absolute log2fold-change > 2. Peaks lacking gene annotations were removed, and duplicate gene annotations were collapsed to a single representative entry per gene symbol for comparison with differentially expressed RNA-seq genes.

### Comparison of Gene Expression in Low versus Normal SOX10 Expressing Cells

A database comprised of RNA sequencing data from patient melanoma was obtained from dbGAP with the accession number phs000933^16^. These data were then sorted to remove any newborn or control melanocytes and filtered by normalized SOX10 read counts. Samples with a normalized SOX10 read count of <200 were categorized as low SOX10 and remaining samples were categorized as normal SOX10. Normalized read counts of target genes were isolated and outliers were identified using the ROUT test with Q set at 1%. The filtered data were then plotted and statistically compared using the Mann-Whitney test for nonparametric data sets in GraphPad Prism.

## Data Availability

All raw and processed RNA-seq files can be accessed in Gene Expression Omnibus (GEO) under accession number GSE335958. All raw and processed ATAC-seq data can be accessed under accession number GSE335959. All uncropped blots, densitometry, luminescence for cell proliferation studies, and raw Cq values can be found in the source data. SOX10 CUT&RUN data were accessed from GEO under accession GSE172069. Patient RNA-seq data was accessed from dbGAP under accession phs000933. All plasmids and cell lines generated in this study will be freely provided upon reasonable request.

## References

1 Siegel, R. L., Kratzer, T. B., Wagle, N. S., Sung, H. & Jemal, A. Cancer statistics, 2026. CA: A Cancer Journal for Clinicians 76, e70043 (2026). 10.3322/caac.70043

2 Shakhova, O. et al. Sox10 promotes the formation and maintenance of giant congenital naevi and melanoma. Nature Cell Biology 14, 882–890 (2012). 10.1038/ncb2535

3 Szumera-Ciećkiewicz, A. et al. SOX10 is as specific as S100 protein in detecting metastases of melanoma in lymph nodes and is recommended for sentinel lymph node assessment. European journal of cancer (1990) 137, 175–182 (2020). 10.1016/j.ejca.2020.06.037

4 Sasaki, K. et al. SOX10 Inhibits T Cell Recognition by Inducing Expression of the Immune Checkpoint Molecule PD-L1 in A375 Melanoma Cells. Anticancer research 43, 1477–1484 (2023). 10.21873/anticanres.16296

5 Yokoyama, S. et al. SOX10 Regulates Melanoma Immunogenicity through an IRF4– IRF1 Axis. Cancer Research 81, 6131–6141 (2021). 10.1158/0008-5472.can-21-2078

6 Rosenbaum, S. R. et al. SOX10 Loss Sensitizes Melanoma Cells to Cytokine-Mediated Inflammatory Cell Death. Molecular cancer research 22, 209–220 (2024). 10.1158/1541-7786.MCR-23-0290

7 Rosenbaum, S. R. et al. SOX10 requirement for melanoma tumor growth is due, in part, to immune-mediated effects. Cell reports (Cambridge) 37, 110085 (2021). 10.1016/j.celrep.2021.110085

8 Takahashi, M. et al. DrugMap: A quantitative pan-cancer analysis of cysteine ligandability. Cell 187, 2536–2556 e2530 (2024). 10.1016/j.cell.2024.03.027

9 Capparelli, C. et al. Targeting SOX10-deficient cells to reduce the dormant-invasive phenotype state in melanoma. Nature communications 13, 1381–1316 (2022). 10.1038/s41467-022-28801-y

10 Wouters, J. et al. Robust gene expression programs underlie recurrent cell states and phenotype switching in melanoma. Nat Cell Biol 22, 986–998 (2020). 10.1038/s41556-020-0547-3

11 Thongpradit, S. et al. Novel SOX10 Mutations in Waardenburg Syndrome: Functional Characterization and Genotype-Phenotype Analysis. Front Genet 11, 589784 (2020). 10.3389/fgene.2020.589784

12 Newsom-Stewart, C. M., Bhatt, D. P., Major, M. B. & Kaufman, C. K. Mapping of the hSOX10 Proximal Protein Interactome in Human Melanoma. Journal of proteome research (2026). 10.1021/acs.jproteome.5c00852

13 Buckley, D. L. et al. HaloPROTACS: Use of Small Molecule PROTACs to Induce Degradation of HaloTag Fusion Proteins. ACS Chem Biol 10, 1831–1837 (2015). 10.1021/acschembio.5b00442

14 Echavarria-Consuegra, L. & Goodfellow, I. Proof of concept: targeted protein degradation of the stress granules component G3BP1 as an antiviral strategy against norovirus infection. Antimicrob Agents Chemother 70, e0111825 (2026). 10.1128/aac.01118-25

15 Zhao, D., Yin, Z., Soellner, M. B. & Martin, B. R. Scribble sub-cellular localization modulates recruitment of YES1 to regulate YAP1 phosphorylation. Cell Chemical Biology 28, 1235–1241.e1235 (2021). 10.1016/j.chembiol.2021.02.019

16 Krauthammer, M. et al. Exome sequencing identifies recurrent mutations in NF1 and RASopathy genes in sun-exposed melanomas. Nat Genet 47, 996–1002 (2015). 10.1038/ng.3361

17 Cao, J. et al. An easy and efficient inducible CRISPR/Cas9 platform with improved specificity for multiple gene targeting. Nucleic Acids Research 44, e149–e149 (2016). 10.1093/nar/gkw660

18 Prasad, M. K. et al. SOX10 directly modulates ERBB3 transcription via an intronic neural crest enhancer. BMC Developmental Biology 11, 40 (2011). 10.1186/1471-213X-11-40

19 Jiao, Z. et al. Direct Interaction of Sox10 with the Promoter of Murine Dopachrome Tautomerase (Dct) and Synergistic Activation of Dct Expression with Mitf. Pigment Cell Research 17, 352–362 (2004). 10.1111/j.1600-0749.2004.00154.x

20 Baggiolini, A. et al. Developmental chromatin programs determine oncogenic competence in melanoma. Science 373, eabc1048 (2021). 10.1126/science.abc1048

21 Hu, W. et al. NAV2 facilitates invasion of cutaneous melanoma cells by targeting SNAI2 through the GSK-3β/β-catenin pathway. Archives of Dermatological Research 311, 399– 410 (2019). 10.1007/s00403-019-01909-w

22 McGrail, K. et al. Transcriptional reprogramming triggered by neonatal UV radiation or Lkb1 loss prevents BRAFV600E-induced growth arrest in melanocytes. Oncogene 44, 1592–1608 (2025). 10.1038/s41388-025-03339-7

23 Tudrej, K. B., Czepielewska, E. & Kozłowska-Wojciechowska, M. SOX10-MITF pathway activity in melanoma cells. Arch Med Sci 13, 1493–1503 (2017). 10.5114/aoms.2016.60655

24 Bondeson, D. P. et al. Systematic profiling of conditional degron tag technologies for target validation studies. Nature Communications 13, 5495 (2022). 10.1038/s41467-022-33246-4

25 Ody, B. K. et al. Synthesis and Evaluation of Cereblon-Recruiting HaloPROTACs. ChemBioChem 24, e202300498 (2023). 10.1002/cbic.202300498

26 Stevens, R. et al. A Systematic Direct-to-Biology Approach Identified Potent Cereblon HaloPROTACs. ACS Chemical Biology 21, 1798–1817 (2026). 10.1021/acschembio.6c00377

27 Bacchiocchi, A. et al. LZTR1 is a melanoma oncogene that promotes invasion and suppresses apoptosis. Oncogene 44, 3974–3984 (2025). 10.1038/s41388-025-03538-2

