## Supplementary Information for "HaloPROTAC-mediated degradation reveals SOX10-dependent gene expression changes in melanoma cells"

* To whom correspondence should be addressed:

**Supplementary Tables S1-S2; Supplementary Figures S1-S8.**

Table S1: Guide RNA (gRNA) and primer sequences.

| | |
|---|---|
| **SOX10 Targeting Single Guide RNA (sgRNA)** | |
| Forward 1 | GGCTCTCCTCTAAACCCCGT |
| Forward 2 | ACTCTTTGCGAGGACCCCGG |
| Reverse 1 | AGTCGTAGCCGCTGAGCACC |
| Reverse 2 | GCTGGCGCCGTTGACGCGCA |
| **Oligonucleotides for sgRNA Cloning** | |
| Forward 1 | ATCTTGTGGAAAGGACGAAACACCGGGCTCTCCTCTAA ACCCCGTGTTTTAGAGCTAGAAATAGCAAGTT |
| Forward 2 | ATCTTGTGGAAAGGACGAAACACCGACTCTTTGCGAGG ACCCCGGGTTTTAGAGCTAGAAATAGCAAGTT |
| Reverse 1 | ATCTTGTGGAAAGGACGAAACACCGAGTCGTAGCCGCT GAGCACCGTTTTAGAGCTAGAAATAGCAAGTT |
| Reverse 2 | ATCTTGTGGAAAGGACGAAACACCGGCTGGCGCCGTTG ACGCGCAGTTTTAGAGCTAGAAATAGCAAGTT |
| **PCR Primers for SOX10 Genotyping** | |
| SOX10 Forward | TGGGACACGGTTTTCCACTT |
| SOX10 Reverse | CTCGGCTACCCTGAATCCAC |
| **RT-qPCR Primers** | |
| ACTIN Forward | AGGCACCAGGGCGTGAT |
| ACTIN Reverse | GCCCACATAGGAATCCTTCTGAC |
| CD200 Forward | ACTGTACTGGAAACGTCACC |
| CD200 Reverse | GCTCTTTCAGGTCCTTTGGA |
| RFTN1 Forward | ACCGACAGATCTCAGAAAACT |
| RFTN1 Reverse | GTGGTCTAAGGAGGAACAGC |
| DKK2 Forward | GCAATAATGGCATCTGTATCCC |
| DKK2 Reverse | GAGTACCATCCAGAGCCG |
| NAV2 Forward | TGGAGCCAAGTACCCAGATGTG |
| NAV2 Reverse | GAGGATTGGAGATGACCACCGA |
| NCKAP5 Forward | TTCCGTGGCTGTGAACAAGTCTAAG |
| NCKAP5 Reverse | AACTGCCCTGTGCTTGTGAATCC |
| TDRD7 Forward | CTAAGGGCTGTCCTGCAGTC |
| TDRD7 Reverse | AGAGTTGCCTTTGGCTTT |
| PKNOX2 Forward | CTTTGTCCACCAGCAGGAACAG |
| PKNOX2 Reverse | GGTGGATTCTCAGGACCTGGAT |
| MSK1 Forward | CTCCTCACTGTCAAGCACGAG |
| MSK1 Reverse | GCCTTTTGAACGATTGTTGCCT |
| CDH1 Forward | GAGTGCCAACTGGACCATTCAGTA |
| CDH1 Reverse | AGTCACCCACCTCTAAGGCCATC |

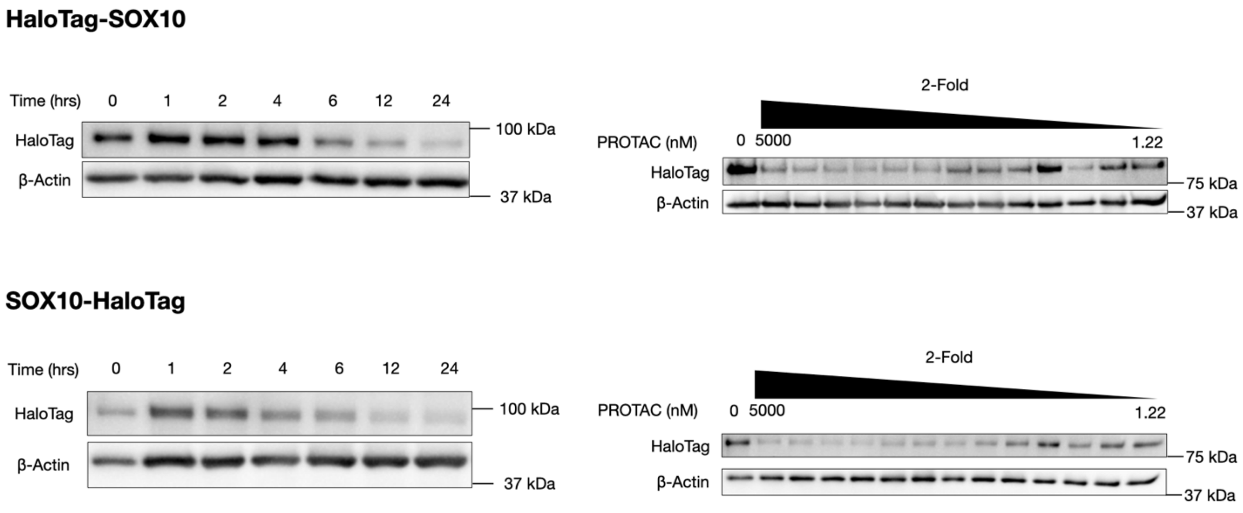


Figure S1: HaloPROTAC3 concentration and time course optimization in primary melanoma cells. SOX10 fusion proteins were stably overexpressed in WW165 cells. Left, Western blot analysis of HaloTag-SOX10 (top) and SOX10-HaloTag (bottom) fusion protein degradation at various time points after treatment with 1.25 μM HaloPROTAC3 (PROTAC) (left). Right, Western blot analysis of HaloTag-SOX10 (top) and SOX10-HaloTag (bottom) fusion protein degradation at varied HaloPROTAC3 concentrations (right). β-actin was used as a loading control.

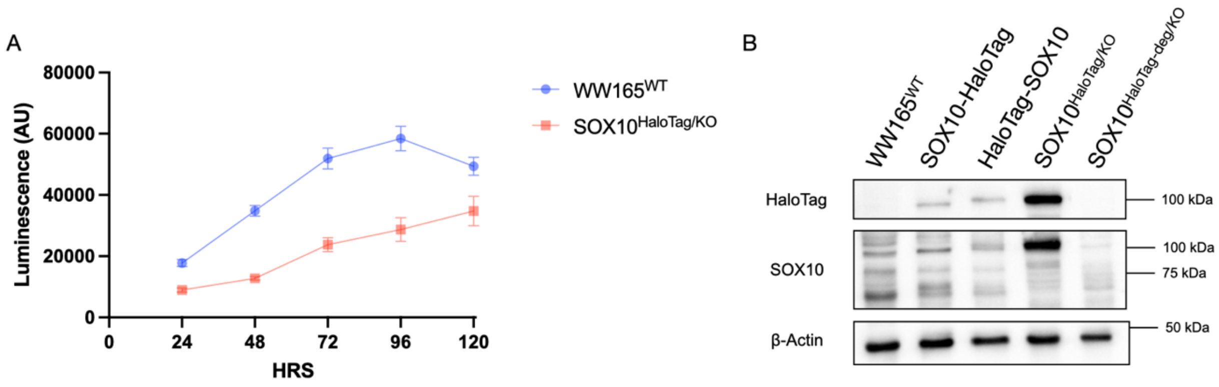


Figure S2: Characterization of SOX10 knockout WW165 melanoma cells expressing SOX10-HaloTag ($SOX10^{HaloTag/KO}$). (A) Growth curves comparing wild-type WW165 ($WW165^{WT}$) and $SOX10^{HaloTag/KO}$. Error bars, standard deviation (n=16). (B) Western bloting of SOX10 fusion protein and endogenous SOX10 expression in cell lines used in this study: from left to right, $WW165^{WT}$, $WW165^{WT}$ expressing SOX10-HaloTag, $WW165^{WT}$ expressing HaloTag-SOX10, $SOX10^{HaloTag/KO}$, and $SOX10^{HaloTag/KO}$ treated with 1.25 μM HaloPROTAC3 for 48 hours ($SOX10^{HaloTag\text{-}deg/KO}$). β-actin, loading control.

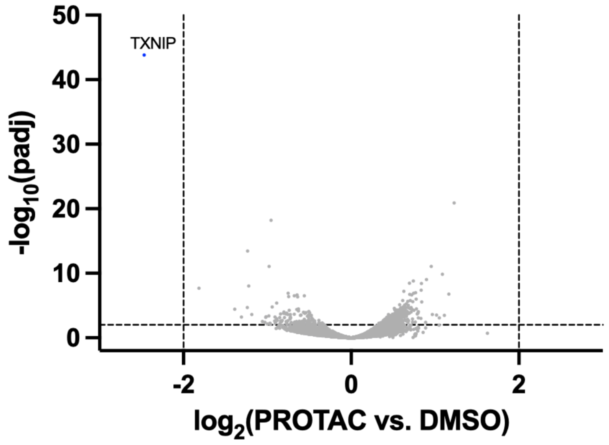


Figure S3: RNA-seq of wild-type WW165 cells treated with HaloPROTAC3. Volcano plot of RNA-seq data from wild-type WW165 cells treated with either DMSO or HaloPROTAC3 (1.25 µM) for 48 hours. P-values, unpaired, two-tailed t-test.

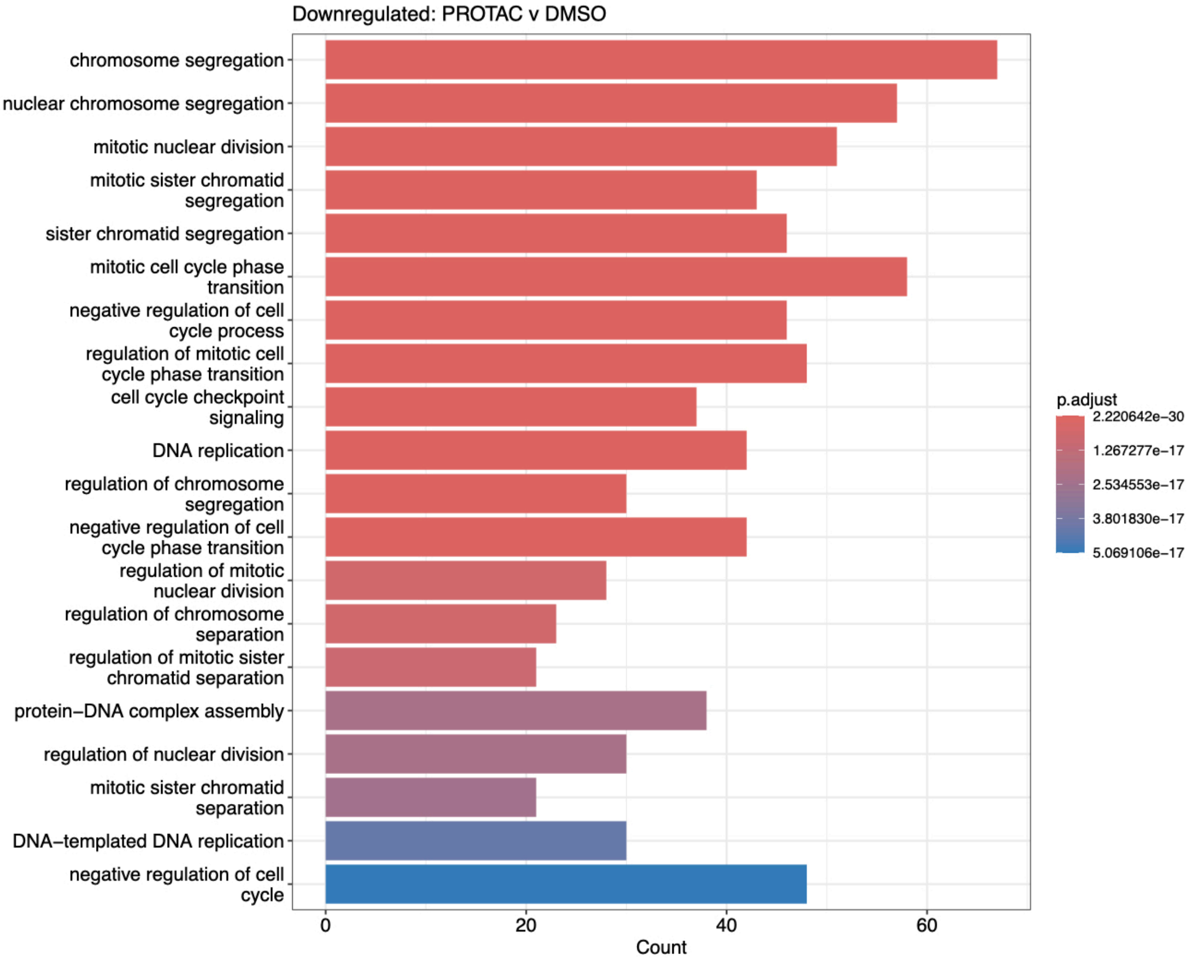


Figure S4: Gene ontology (GO) biological process analysis of differentially expressed genes after SOX10 targeted degradation in SOX10$^{HaloTag/KO}$ cells. Significantly downregulated genes were analyzed with enrichGO Function from the clusterProfiler R package and assigned to respective biological processes. Significance was determined as $\log_2$(PROTAC/DMSO) $\leq -2$ with a $p_{adjusted}$ value of $\leq 0.01$ for each dataset.

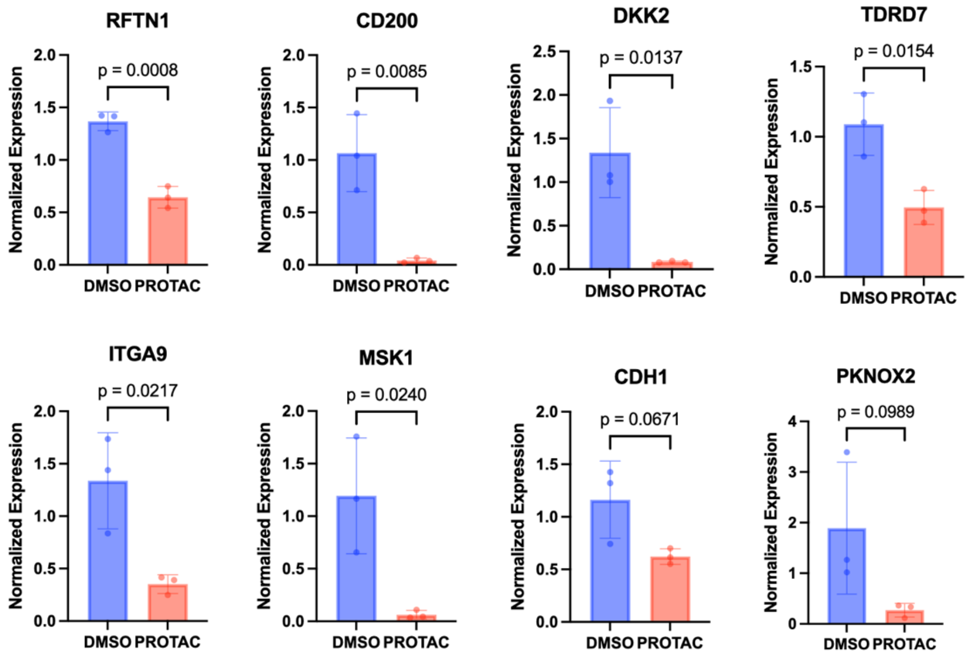


Figure S5. Changes in transcript levels upon SOX10 degradation. RT-qPCR with specific primers for indicated genes was performed on mRNA isolated from SOX10$^{HaloTag/KO}$ cells treated with DMSO control (blue) or HaloPROTAC3 (PROTAC, red) for 48 hours. P-value, unpaired t-test; error bars, standard deviation. Each data point represents one biological replicate averaged from three technical replicates.

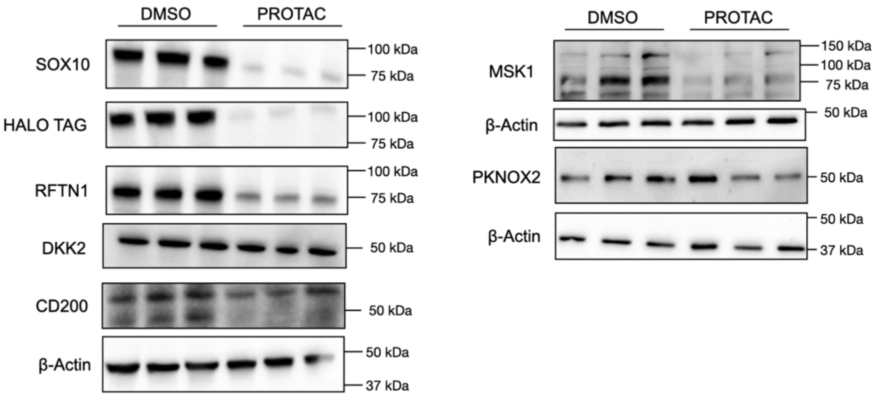


Figure S6. Changes in protein levels upon SOX10 degradation. Western blot assessment of protein expression of SOX10$^{HaloTag/KO}$ cells treated with DMSO or HaloPROTAC3 (PROTAC) for 48 hours. Each sample represents one biological replicate. β-actin, loading control.

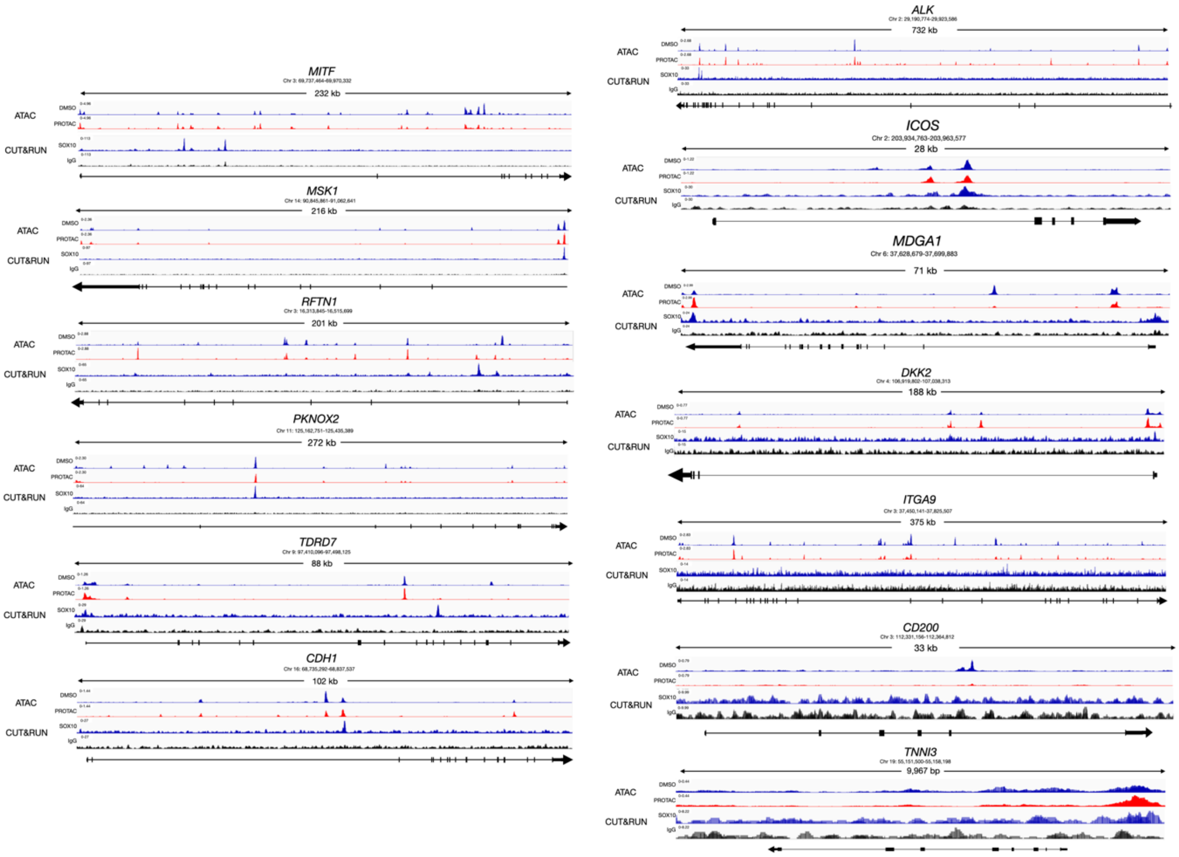


Figure S7. Chromatin accessibility after SOX10 degradation and direct binding of novel target genes by SOX10. Top two tracks, ATAC-seq signal from SOX10$^{HaloTag/KO}$ cells treated with DMSO (blue) or HaloPROTAC3 (red) for 48 hours prior to sequencing. Signal represents the average of 3 biological replicates. Bottom two tracks, reanalysis of SOX10 CUT&RUN (blue) vs. IgG control (black) at the locus of interest[1]. Each track represents the average of 3 replicates. *MITF*, positive control; *TNNI3*, negative control not expressed in skin; other genes are candidate SOX10 targets. Tracks are arranged in order of descending relative intensities from top to bottom. A representative diagram of the gene body is displayed below each track with black boxes representing exons and arrow showing directionality.

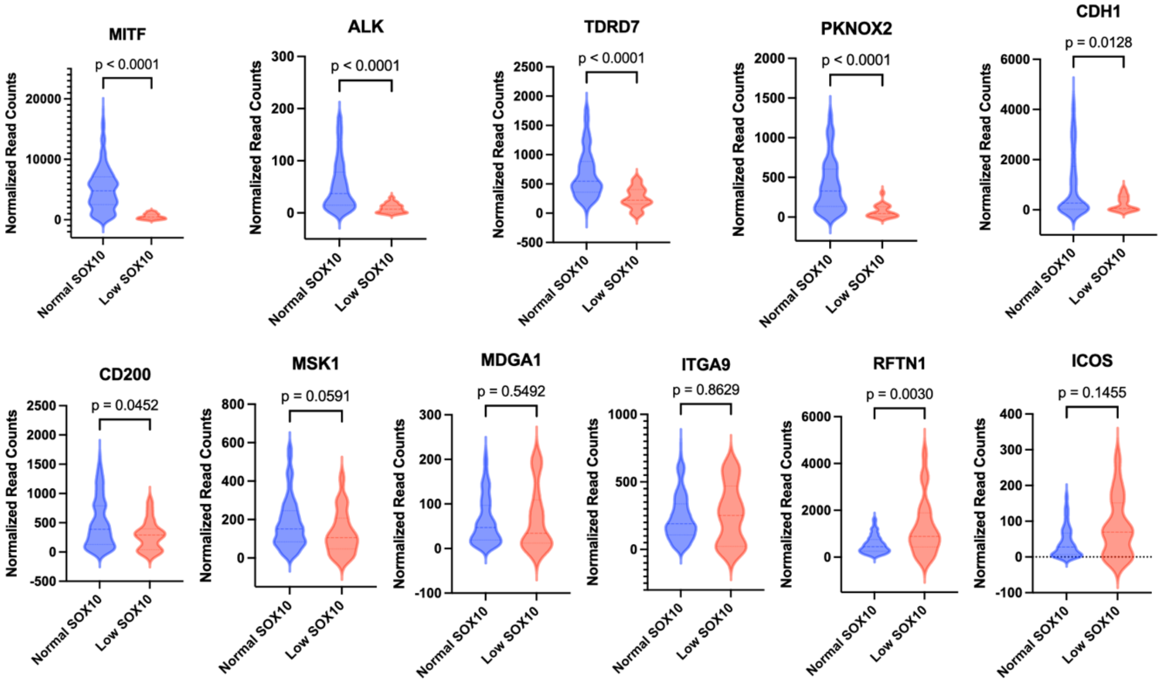


Figure S8. Correlation of novel genes of interest with SOX10 levels in melanoma. Melanoma patient RNA-seq data from Krauthammer et al.[2] were reanalyzed to determine expression levels of novel candidate SOX10 target genes in low- vs. high-SOX10 expressing melanomas. Normalized read counts comparing specific gene expression in low-*SOX10* expressing melanomas (normalized *SOX10* read count < 200, red; n = 22) versus high-*SOX10* expressing melanomas (normalized *SOX10* read count > 200, blue; n = 275). A ROUT test was performed to remove statistically significant outliers (Q=1%). P-value, Mann-Whitney test.

| Gene Name | RNA-seq | ATAC-seq | RT-qPCR | Western Blot | SOX10 CUT&RUN | Patient RNA-seq |
|---|---|---|---|---|---|---|
| NCKAP5 | green | green | green | green | green | green |
| NAV2 | green | green | green | green | green | green |
| TDRD7 | green | green | green | gray | green | green |
| MSK1 | green | green | green | green | green | orange |
| CHD1 | green | green | orange | gray | green | green |
| ALK | green | green | gray | gray | green | green |
| PKNOX2 | green | green | orange | orange | green | green |
| ITGA9 | green | green | green | gray | orange | orange |
| CD200 | green | green | green | green | orange | green |
| RFTN1 | green | green | green | green | green | orange |
| DKK2 | green | green | green | orange | orange | gray |
| MDGA1 | green | green | gray | gray | green | orange |
| ICOS | green | green | gray | gray | green | orange |

Table S2: Summary of evidence for novel SOX10 target genes. RNA-seq: green indicates statistically significant downregulation when $SOX10^{HaloTag/KO}$ cells were treated with HaloPROTAC3 compared to DMSO for 48 hours. ATAC-seq: green indicates statistically significant decrease in chromatin accessibility when $SOX10^{HaloTag/KO}$ cells were treated with HaloPROTAC3 compared to DMSO for 48 hours. RT-qPCR: green indicates statistically significant ($p < 0.05$, Student's t-test) RNA-level downregulation after SOX10-HaloTag degradation; orange indicates no statistically significant downregulation after SOX10 degradation; gray indicates no amplification. Western blot: green indicates protein-level downregulation after SOX10 degradation; orange indicates no protein-level downregulation; gray indicates no detection with reported antibody or no antibody available. SOX10 CUT&RUN: green indicates specific SOX10 CUT&RUN signal in promoter or gene body; orange indicates no SOX10 binding above IgG background. Patient RNA-seq: green indicates statistically significant downregulation in low- vs. high-SOX10-expressing melanoma patient cells or tumors ($p < 0.05$, nonparametric Mann-Whitney test after the data was filtered for outliers using the ROUT test, Q=1%); orange indicates no statistically significant downregulation; gray indicates no detected expression.

**References**


1 Baggiolini, A. *et al.* Developmental chromatin programs determine oncogenic competence in melanoma. *Science* **373**, eabc1048 (2021). https://doi.org/10.1126/science.abc1048

2 Krauthammer, M. *et al.* Exome sequencing identifies recurrent mutations in NF1 and RASopathy genes in sun-exposed melanomas. *Nat Genet* **47**, 996–1002 (2015). https://doi.org/10.1038/ng.3361
